# Early life low-calorie sweetener consumption impacts energy balance during adulthood

**DOI:** 10.1101/2022.10.23.513433

**Authors:** Anna M.R. Hayes, Linda Tsan, Alicia E. Kao, Grace M. Schwartz, Léa Décarie-Spain, Logan Tierno Lauer, Molly E. Klug, Lindsey A. Schier, Scott E. Kanoski

**Author notes:** These authors contributed equally (Anna M.R. Hayes, Linda Tsan). Correspondence: Scott Kanoski, Current Address: 3616 Trousdale Parkway, AHF-252, Los Angeles, CA 90089-0372.

## Abstract

Children frequently consume beverages sweetened with either sugars (sugar-sweetened beverages; SSB) or low-calorie sweeteners (LCS). Here we evaluated the effects of habitual early life consumption of either SSB or LCS on energy balance later during adulthood. Male and female rats were provided with chow, water, and a solution containing either SSB (sucrose), LCS [acesulfame potassium (ACE-K) or stevia], or control (no solution) during the juvenile and adolescent periods (postnatal days 26-70). SSB or LCS consumption was voluntary and restricted within federal recommended daily limits. When subsequently maintained on a cafeteria-style junk food diet (CAF; various high-fat, high-sugar foods) during adulthood, ACE-K-exposed rats demonstrated reduced caloric consumption vs. controls, which contributed to lower body weights in female but not male ACE-K rats. These discrepant intake and body weight effects in male ACE-K rats are likely based on reduced gene expression of thermogenic indicators (*UCP1, BMP8B*) in brown adipose tissue. Female stevia-exposed rats did not differ from controls in caloric intake or body weight, yet they consumed more SSB during adult CAF exposure. No SSB-exposed rats, neither male nor female, differed from controls in adult total caloric consumption or body weight measures. Collective results reveal that early life LCS consumption alters sugar preference, body weight, and gene expression for markers of thermogenesis during adulthood, with both sex- and sweetener-dependent effects.

## 1. Introduction

Early life consumption of Western diet is associated with increased caloric intake and susceptibility to obesity and metabolic dysfunction during adulthood [1–3]. The underlying nutritive mechanisms mediating these effects, however, are poorly understood given that established rodent obesogenic diet models vary from healthy control diets in two primary ways: [1] elevated dietary fat (particularly saturated fatty acids), and [2] elevated sugar. Given that children consume the most sugar of any age group, with sugar-sweetened beverages (SSBs) constituting a major source of sugar in their diet [4,5], it is imperative to understand the long-lasting implications of habitual early life sugar consumption, without concomitant elevated dietary fat, on eating patterns and body weight regulation during adulthood.

Based on the surging increase in obesity and nutrition-related co-morbidities among children and adolescents [6,7], increased focus has been placed on dietary and lifestyle interventions to better support healthy growth trajectories in early life [8,9]. One such strategy is to reduce sugar consumption. Decreasing sugar consumption while preserving pleasurable sweet taste in the diet can at least theoretically be achieved by consuming foods and beverages in which sugars have been fully or partially replaced with low-calorie sweeteners (LCSs). However, evidence from both humans and rodent models is mixed and challenges the efficacy of using LCSs for weight management and energy balance maintenance [10–13]. Furthermore, LCS consumption has been shown to alter glucose homeostasis and gut peptide release in humans [14,15] and influence cephalic phase physiological signaling in rodent models [11,16]. Given that LCS consumption in children increased by ~200% from 1999 to 2012 [17], thus representing a societal shift towards caloric sugar replacement with LCSs during critical stages of development, it is important to mechanistically understand the long-term impacts of early life LCS consumption on subsequent energy balance parameters during adulthood.

LCSs and caloric sugars bind to the same sweet receptors [18], yet unlike sugars, LCSs provide minimal-to-no energy to the body and comprise a wide variety of chemical structures [19]. Acesulfame potassium (ACE-K) is a commonly used synthetic LCS that retains its sweetness at high temperatures [20,21], thus making it an attractive LCS for use in a variety of processed and baked foods. Stevia, or the collection of Steviol glycosides, is extracted from the leaves of *Stevia rebaudiana* Bertoni and is one of the most popular ‘natural’ sweeteners with recent exponential increase in its use in food and beverage products [20,22]. Although a number of studies have examined the effects of caloric sugars or LCSs on energy balance control [10–13,23], we emphasize two primary gaps in the literature: [1] the majority of mechanistic physiological rodent model research has used the LCS saccharin [e.g., [24–29]], which is not commonly consumed by humans, and [2] rodent model research often involves sugar or LCS consumption that is far in excess of what is consumed in human populations and involves involuntary consumption conditions where sugar or LCS is supplemented in drinking water. Therefore, in the present study we aimed to fill critical gaps in the literature by assessing the long-term impacts of early life consumption of sugar or LCSs commonly consumed by humans (ACE-K or stevia) on eating behavior, metabolic outcomes, and body weight regulation during adulthood using a rat model where consumption is voluntary and restricted within federal recommended daily limits. To evaluate potential impacts of early life sugar or LCS on food choice during adulthood, our adult diet model employed an obesogenic cafeteria-style junk food diet (CAF) that involved a variety of food options high in dietary fat, sugar, or both.

## 2. Methods

### 2.1 Study design

Male and female Sprague Dawley rats [postnatal day (PN) 25; 50-70g] were purchased from Envigo (Indianapolis, IN, USA). Upon arrival, juvenile rats were housed individually in a climate controlled (22–24 °C) environment with a 12:12 hour light/dark cycle (lights off at 18:00) and maintained on *ad libitum* standard rodent chow (Lab Diet 5001; PMI Nutrition International, Brentwood, MO, USA; 29.8% kcal from protein, 13.4% kcal from fat, 56.7% kcal from carbohydrate) and water. From PN 26-70, a period that spans juvenile, adolescence, and young adulthood [30], rats received their experimental early life diets (section 2.2) following randomization into groups of comparable weights. Body weights were recorded daily whereas water consumption and chow intake were recorded 3 times per week. All procedures were approved by the Institutional Animal Care and Use Committee at the University of Southern California.

### 2.2 Early Life Diets

Juvenile male and female rats (n=9 per sex) were given daily amounts of acesulfame potassium (ACE-K group; catalog # A2815, Spectrum Chemical, Gardena, CA, USA; 0.1% ACE-K weight/volume (w/v) in reverse osmosis (RO) water; ~15 mg/kg), sucrose (10% KCAL SUG group; C&H Crockett, CA, USA; 11% w/v sucrose in RO water), or stevia (STEVIA group; JG Group, Ontario, Canada; 0.033% stevia w/v in RO water; ~4 mg/kg). The dose of ACE-K and stevia delivered daily was calculated to be within U.S. FDA’s acceptable daily intake (ADI) of each sweetener by body weight. Throughout the early life diet exposure period, the ACE-K and stevia concentrations were fixed, but the volumes administered were increased daily and individually for each rat to provide a fixed mg/kg dose across growth. LCSs were administered for voluntary consumption via a syringe into empty rodent sippers (part #ST7.0-BO with standard stoppers manufactured by Alternative Design Manufacturing & Supply, Siloam Springs, AR, USA). Each sipper had a vinyl cap (6mm inner diameter, Jocon SF9000) attached to its end to prevent leakage and was placed on the wire rack adjacent to the rat’s chow and water bottle. The volume of sucrose delivered daily was based on general guidelines for sugar intake (<10% of total calories per day; Dietary Guidelines for Americans [31]) and was calculated to be equivalent to approximately 10% of the calories consumed from chow the previous day for each individual rat. Sucrose was provided using a syringe to inject the exact volume into a 100 ml glass bottle containing a #6.5R rubber stopper and OT-100 straight 2.5” sipper tube (Ancare, Bellmore, NY, USA). Voluntary consumption of the entire rations of ACE-K, sucrose, and stevia was verified daily by inspecting the tube for all animals. Each experimental group had its own control group (CTL; n=9 per sex) that was provided a tube filled with RO water at an equivalent volume as the experimental groups. The concentrations of ACE-K (0.1% w/v) and stevia (0.033% w/v) were determined based on our preliminary two bottle preference tests (concentrations preferred to water; data not shown) as well as concentrations used in published studies [32,33]. The concentration of sucrose (11% w/v) matched the concentration used in our previous studies [34–37], which was chosen to be comparable to the concentrations found in sugar-sweetened beverages commonly consumed by humans in modern Western cultures.

### 2.3 Glucose Tolerance Test (GTT)

Rats were habituated to oral gavage, food-restricted 22 h prior to the GTT as previously described [38], and water restricted at the start of the dark cycle (15 h prior to the GTT). Five minutes prior to oral gavage with dextrose, baseline blood glucose readings were obtained from the tail tip and recorded by a blood glucose meter (One-touch Ultra2, LifeScan Inc., Milpitas, CA, USA). Each animal was then orally gavaged with dextrose in 0.9% sterile saline (20% w/v, 9.5 ml/kg by body weight) and tail tip blood glucose readings were obtained at 5, 10, 30, 60 and 120 min after gavage.

### 2.4 Cafeteria Diet Access in Adulthood

At PN 82 (in adulthood), rats were transferred to hanging wire-bottom cages and were provided free access to a cafeteria (CAF) diet consisting of potato chips (Ruffles), chocolate peanut butter cups (Reese’s minis), 45% kcal high fat/sucrose chow (Research Diets D12451, New Brunswick, NJ, USA), a bottle of 11% weight/volume (w/v) high fructose corn syrup (HFCS)-55 solution (Best Flavors, Orange, CA, USA), and water. The concentration of sugar (11% w/v) was based on prior studies to model commonly consumed SSBs in humans [30,39]. In each home cage, there were individual, designated food hoppers for accurate measurement of each solid food type. The potato chips and chocolate peanut butter cups were crushed or chopped to facilitate consumption through the hoppers. Papers were placed underneath the hanging wire cages to collect and weigh food spillage. Body weights and food intake were measured 3x/week during CAF diet access.

### 2.5 Open Field

At PN 114, while on the CAF diet, the rats were tested in the open field task, a behavioral paradigm used in rodents to evaluate innate anxiety-like and locomotor activity behaviors [40]. Open field procedures are derived from Suarez et al. [41] and testing occurred during the light cycle. The apparatus consisted of a gray arena (60 cm *×* 56 cm), with a designated center zone within the arena (19 cm *×* 17.5 cm), placed under diffused lighting (30 lux). Animals were placed in the center of the box and allowed to freely explore for 10 minutes. The apparatus was cleaned with 10% ethanol between rats. Anxiety-like behavior was measured as time spent in the center zone, and locomotor activity was measured as distance travelled (m) in the apparatus during the task as quantified by a video tracking system (ANY-maze, Stoelting Co., Wood Dale, IL, USA).

### 2.6 Quantitative polymerase chain reaction (qPCR)

Following injection of an anesthetic (ketamine 90 mg/kg, xylazine 2.7 mg/kg, and acepromazine 0.64 mg/kg), interscapular brown adipose tissue (BAT) was dissected and flash frozen in a beaker filled with isopentane and surrounded by dry ice. To quantify the relative *UCP1* and *BMP8B* mRNA expression between the control (n=18, 9 females and 9 males) and ACE-K (n=18, 9 females and 9 males) rats in the BAT, reverse transcriptase quantitative polymerase chain reaction (RT-qPCR) was performed as previously described [42]. Briefly, total RNA was extracted from each sample using the RNeasy Lipid Tissue Mini Kit (Cat No. 74804, Qiagen, Hilden, Germany) according to the manufacturer’s instructions. Total RNA concentration per sample was measured using a NanoDrop Spectrophotometer (ND-ONE-W, ThermoFisher Scientific, Waltham, MA, USA). RNA (2000 ng) was reverse transcribed to cDNA using the QuantiTect Reverse Transcription Kit (Cat No. 205311, Qiagen) according to the manufacturer’s instructions. The following probes for rat were utilized from Applied Biosystems: *Rplp0* (Rn03302271_gH), *Ucp1* (Rn00562126_m1) and *Bmp8b* (Rn01516089_gH). qPCR was performed with Applied Biosystems TaqMan Fast Advanced Master Mix (Cat#4444557, ThermoFisher Scientific) using the Applied Biosystems QuantStudio 5 Real-Time PCR System (ThermoFisher Scientific). All reactions were run in triplicate, and control wells without cDNA template were included to verify the absence of genomic DNA contamination. The triplicate Ct values for each sample were averaged and normalized to *Rplp0* expression. The comparative 2-ΔΔCt method was utilized to quantify relative expression levels between groups for the genes of interest.

### 2.7 Statistical analysis

Data are presented in all figures as means ± standard error of the mean (SEM) for error bars. ACE-K, sugar, and stevia groups were analyzed with their respective control (but not compared to each other) because these groups were evaluated in separate experiments. Analyses were performed using Prism software (GraphPad Inc., version 8.4.2), with significance considered at *p* < 0.05. Two-way analysis of variance (ANOVA) was used to assess total caloric intake of standard chow, total caloric intake of CAF diet, and body weight, employing Sidak’s multiple comparisons test when significant main effects were indicated. For caloric intake of standard chow, total kcals consumed was assessed with factors of group (between-subjects), sex (between-subjects), and with the group *×* sex interaction incorporated into the model. For caloric intake of CAF diet, analyses were performed for total kcals [group (between-subjects), sex (between-subjects), group *×* sex interaction] as well as kcals consumed over time per sex [group (between-subjects), time (within-subjects), group *×* time interaction]. Because significant main effects for sex were indicated for caloric intake of standard chow and CAF diet, separate analyses were performed for males and females for the other outcomes. Percentage caloric consumption of each of the CAF dietary components over time was assessed separately for males and females by either repeated measures ANOVA or mixed effects analyses with group (between-subjects, fixed), time (within-subjects, fixed), and the group *×* time interaction as factors. Repeated measures ANOVA was performed for body weight analyses with group (between-subjects), time (within-subjects), and the group *×* time interaction as factors in the model. A mixed-effects analysis was used for glucose tolerance, with fixed effects of group, time, and the group *×* time interaction in the model and subject set as a random effect (making this a mixed-effects analysis). A two-tailed unpaired t-test was used to analyze outcomes from the open field behavioral task (anxiety-like behavior, locomotor activity) as well as mRNA expression of *BMP8B* and *UCP1*.

## 3. Results

### 3.1 Early life habitual ACE-K consumption reduces total caloric intake of a cafeteria diet during adulthood

During the ACE-K access period with maintenance on standard chow and water (PN 26-70; timeline shown in Fig. 1A), neither body weights (Fig. 1B-C), chow intake (Fig. 1D-F), nor water consumption (Supplementary Fig. S1) significantly differed between groups for either sex. At PN 75, after a brief washout period without ACE-K access, there were no differences in glucose tolerance between groups for either sex (Fig. 1G-H).

**Figure 1:**
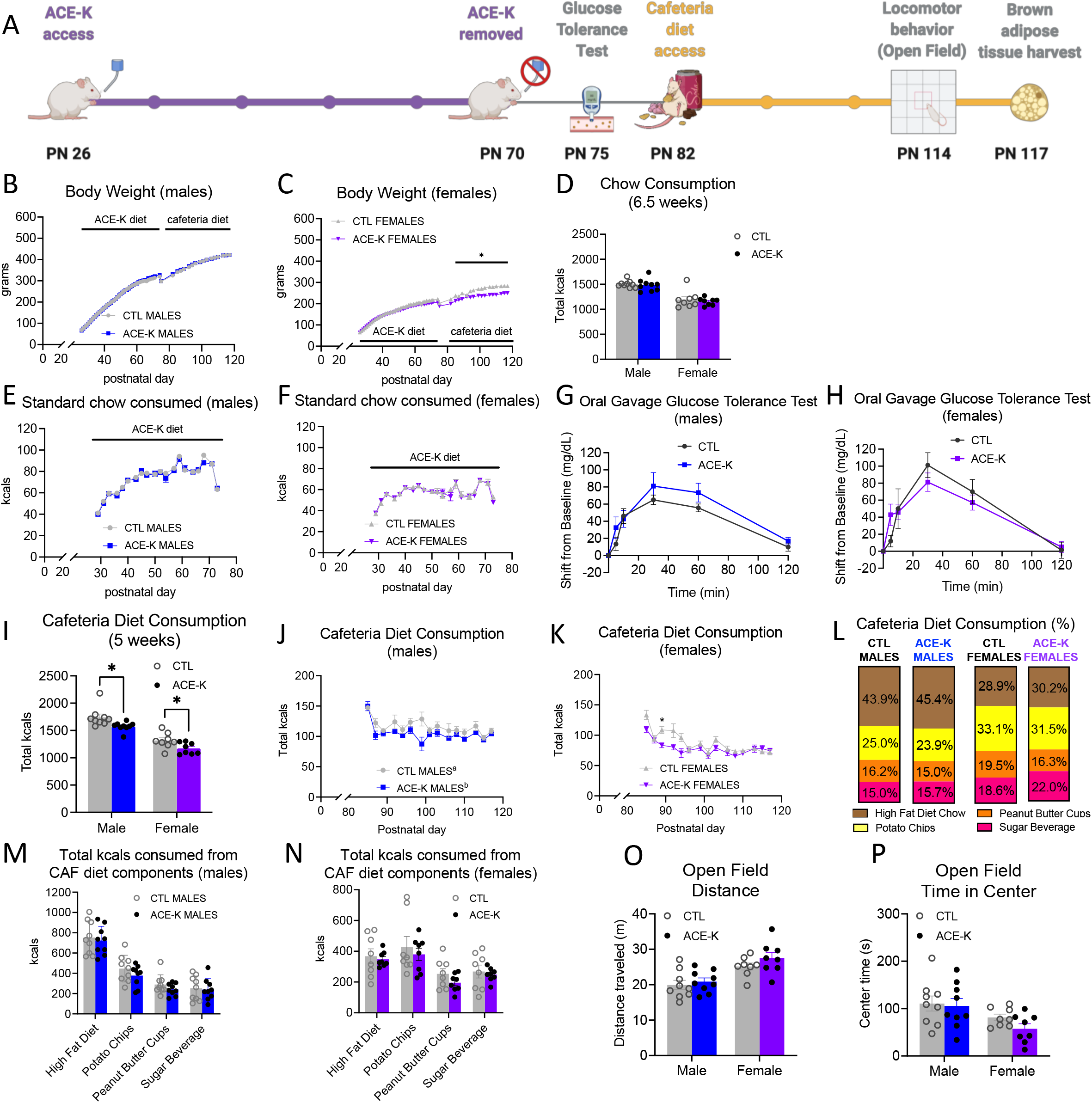
Effects of early life consumption of ACE-K on energy balance parameters during adulthood. Early life ACE-K consumption at the ADI (15 mg/kg; A) had no effect on body weight (B-C), standard chow intake (D-F), or glucose tolerance (G-H) relative to controls that did not have access to ACE-K. When challenged with a cafeteria diet in adulthood, both ACE-K males and females reduced their total caloric intake relative to controls (I-K) with no differences in kcals consumed from the cafeteria diet components (L-N). While reduced caloric intake decreased body weight in females (C), there were no differences in body weight in males (B). These results were not confounded by differences in locomotor activity (O) or anxiety-like behavior (P). Data are represented as mean ± SEM, \**p* < 0.05; PN: postnatal day; CAF: cafeteria; CTL: control; ACE-K: acesulfame potassium; kcal: kilocalories.

During the CAF access period (PN 82-117), male ACE-K rats had similar body weights compared to controls (Fig. 1B), whereas female rats that consumed ACE-K during early life displayed a significant reduction in body weight (*p* < 0.05, Fig. 1C). Both male and female rats previously exposed to ACE-K had significantly reduced overall caloric intake while maintained on the CAF diet (*p* < 0.05, Fig. 1I-K). The overall reduction in calories consumed from the CAF diet over 5 weeks was not driven by differences in consumption of a particular component of the CAF diet (HFD, potato chips, peanut butter cups, and sugar beverage; Fig. 1L-N and Supplementary Figs. S1 and S4), and there were no differences in macronutrient percentages consumed between groups (% kcals from fat, carbohydrate, protein; Supplementary Fig. S4). Neither locomotor activity nor time spent in the center zone of the open field apparatus differed by group (Fig. 1O-P). Collectively, results show that early life ACE-K consumption reduces caloric intake on a CAF diet during adulthood in both sexes, yet this caloric reduction was only accompanied by a reduction in body weight in female ACE-K rats.

### 3.2 Early life habitual sugar consumption does not impact cafeteria diet consumption in adulthood

Early life access to a bottle of sucrose with a caloric content equivalent to 10% of total calories consumed from chow the day before (from PN 26-70; timeline shown in Fig. 2A) did not influence body weight relative to controls (Fig. 2B-C). There were also no group differences in intake of chow (Fig. 2D-F). Water intake decreased in male rats (significant main effect of group and the group *×* time interaction with significant post hoc comparisons at PN 29 and 36, *p* < 0.05; Supplementary Fig. S2) and female rats (significant group *×* time interaction, *p* < 0.05 no significant post hoc comparisons; Supplementary Fig. S2). These differences in water intake were likely due to compensation for the volume of sucrose beverage provided per day (10-20 mL). At PN 75, after daily sucrose access ceased, there were no group differences in glucose tolerance (Fig. 2G-H). The early life sucrose consumption also did not affect adulthood total caloric consumption of a cafeteria diet, nor was consumption of any of the individual dietary components impacted by group in either males or females (Fig. 2I-N and Supplementary Figs. S2 and S4). Likewise, locomotor activity (Fig. 2O) and anxiety-like behavior (Fig. 2P) were unaffected in the 10% KCAL SUG group.

**Figure 2:**
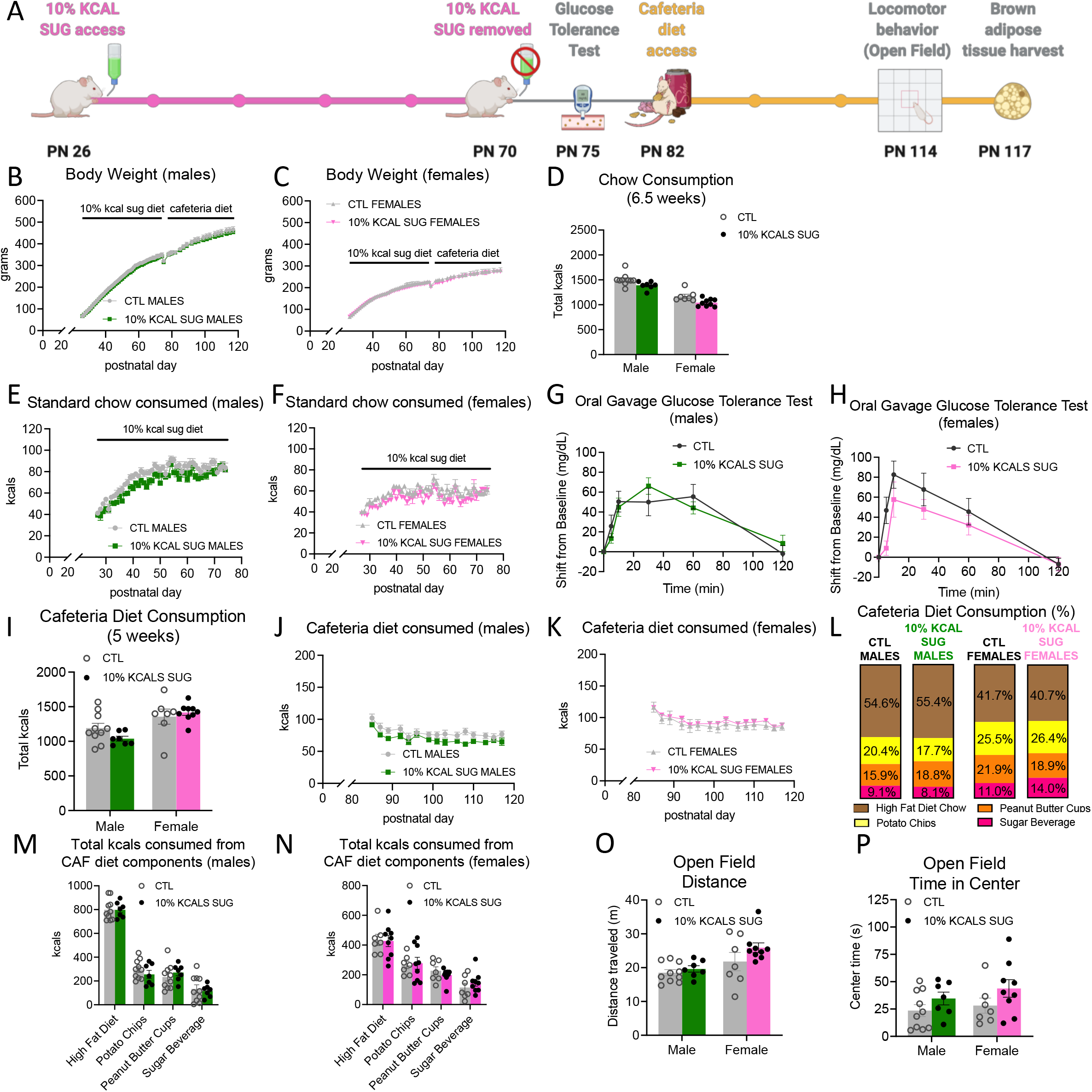
Effects of early life consumption of sugar at 10% of total calories on energy balance parameters during adulthood. Limiting consumption of sugar to 10% of total calories consumed during adolescence (A) had no effect on body weight (B-C), chow intake (D-F), glucose tolerance (G-H), cafeteria diet consumption (I-N), locomotor activity (O), or anxiety-like behavior (P) relative to controls that did not have sugar access during adolescence. Data are represented as mean ± SEM; PN: postnatal day; CAF: cafeteria; CTL: control; SUG: sugar; kcal: kilocalories.

### 3.3 Early life stevia consumption increases sugar-sweetened beverage consumption during adult cafeteria diet exposure

With daily stevia access from PN 26-70 dosed within the federal ADI levels (timeline shown in Fig. 3A), body weights were not significantly different from controls (Fig. 3B-C). Neither males nor females in the stevia group displayed differences in intake of chow (Fig. 3D-F) or water (Supplementary Fig. S3) relative to controls. At PN 75, following cessation of daily stevia access, there were no group differences in glucose tolerance (Fig. 3G-H). Although overall total caloric consumption in stevia-exposed males and females (Fig. 3I-K) was not significantly different relative to controls, stevia-exposed females consumed significantly more of the sugar beverage relative to control females (~16% of total kcals consumed for STEVIA females vs. ~11% of total kcals consumed for CTL females; *p* < 0.05; Fig. 3L, 3N, Supplementary Fig. S3). Despite this increase in calories consumed from the sugar beverage, stevia-exposed female rats did not consume significantly more calories from carbohydrates relative to controls (*p* = 0.11, Supplementary Fig. S3). Consumption of other components of the cafeteria diet (HFD, potato chips, peanut butter cups) was not influenced by early life stevia exposure in females (Fig. 3N and Supplementary Figs. S3-4), and no differences in consumption of the cafeteria diet components were observed for stevia males (Fig. 3L-M and Supplementary Figs. S3-4). Similarly, locomotor activity (Fig. 3O) and anxiety-like behavior (Fig. 3P) were unaffected by early life stevia consumption followed by cafeteria diet consumption during adulthood.

**Figure 3:**
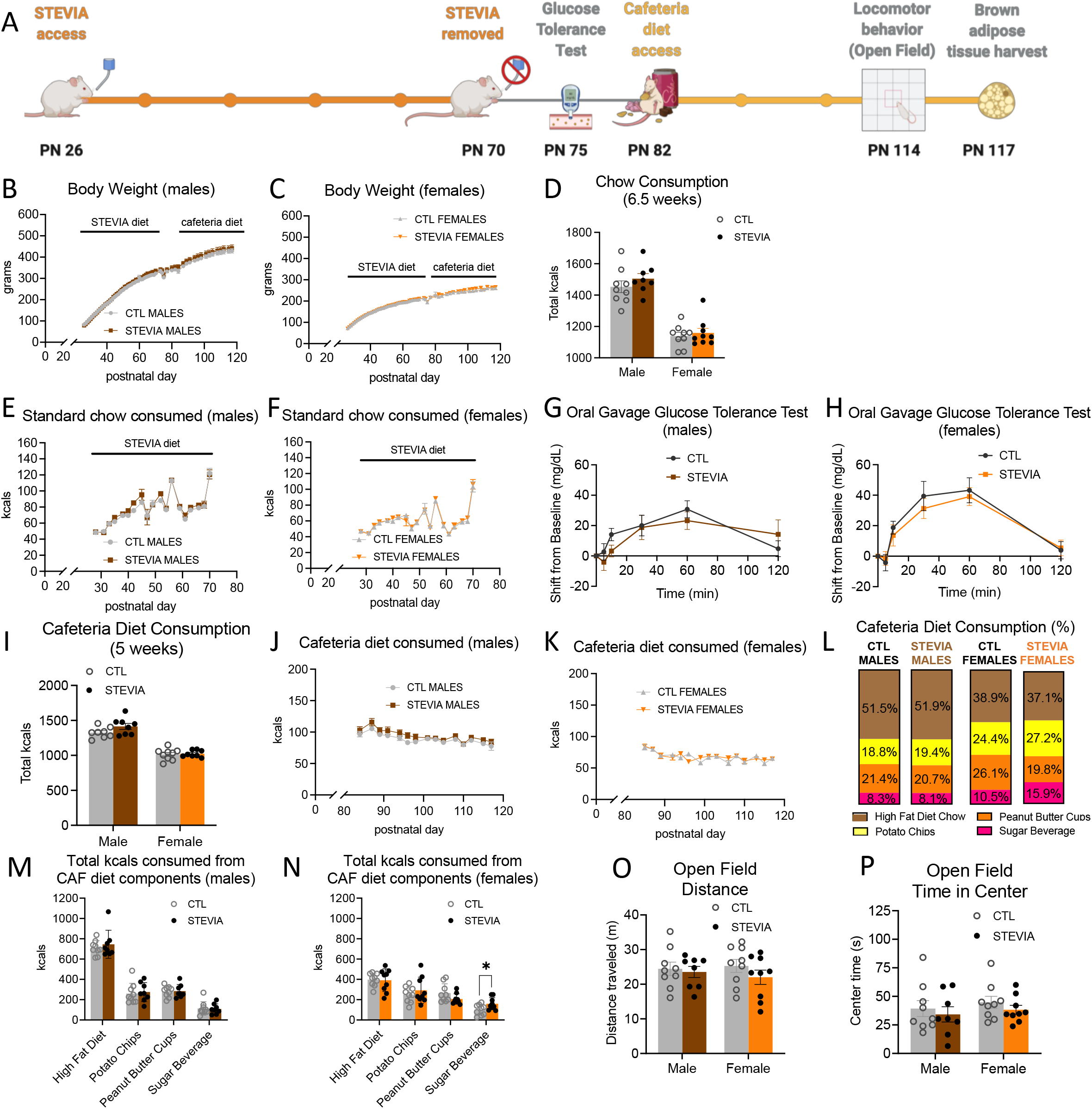
Effects of early life consumption of stevia in rats on energy balance parameters during adulthood. Early life stevia consumption at the ADI (4mg/kg; A) had no effect on body weight (B-C), chow intake (D-F), glucose tolerance (I-J), total calories consumed during adult exposure to cafeteria diet (I-K) relative to controls that did not have sugar access during adolescence. There were no significant differences in percentage of calories consumed from the four cafeteria diet components in either male or female rats (L). Early life stevia consumption did not alter total calories consumed from the cafeteria diet components in male rats during adulthood (M), but it did significantly increase calories consumed of the sugar beverage in female rats during cafeteria diet consumption during adulthood – in the absence of differences among the other three cafeteria diet components (N). Locomotor activity (O) and anxiety like behavior (P) were not influenced by early life stevia exposure. Data are represented as mean ± SEM, \**p* < 0.05; PN: postnatal day; CAF: cafeteria; CTL: control; kcal: kilocalories.

### 3.4 ACE-K consumption during early life reduces brown adipose tissue gene expression of the thermogenic markers, *BMP8B* and *UCP1*, in male rats on a cafeteria diet

To further understand why early life ACE-K exposure reduces consumption of a cafeteria diet in adulthood without a corresponding decrease in body weight in male rats (but not female rats, where both caloric intake and body weighs were reduced), the interscapular BAT of ACE-K rats was analyzed for mRNA expression of *BMP8B* and *UCP1*, two common markers of BAT thermogenesis [43,44]. mRNA levels of *BMP8B* were significantly reduced in ACE-K male rats relative to controls (*p* < 0.05, Fig. 4A), but were not significantly affected in ACE-K female rats (Fig. 4B). Similarly, mRNA levels of *UCP1* were significantly reduced in ACE-K males relative to control rats (*p* < 0.05, Fig. 4C), but were not significantly affected in ACE-K female rats (Fig. 4D). No differences in relative BAT weight (as percentage of body weight) were observed for either male or female ACE-K rats (Fig. 4E-F). Thus, the discrepancy between caloric intake and body weight in male ACE-K rats vs. their controls could be mediated by blunted thermogenesis and subsequent reduced energy expenditure.

**Figure 4:**
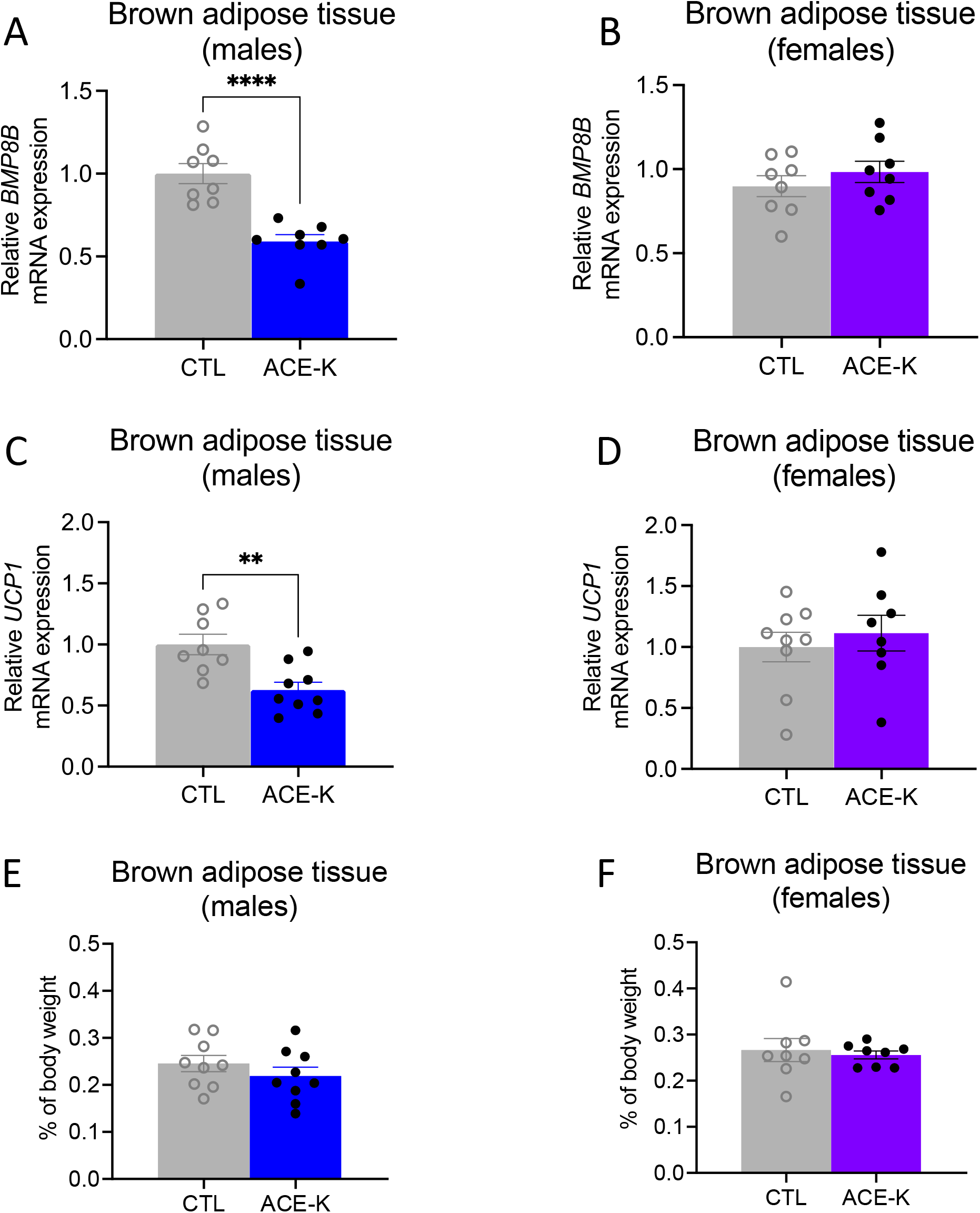
Effect of early life ACE-K on adult brown adipose tissue (BAT) thermogenic markers. Early life ACE-K consumption at the ADI (15mg/kg) decreased relative mRNA expression of *BMP8B* and *UCP1* in male rats (A, C) relative to controls but had no effect in female rats (B, D). These effects were observed in the absence of differences in BAT percentage of total body weight for both sexes (E-F). Data are represented as mean ± SEM, \*\**p* < 0.01, \*\*\*\**p* < 0.0001; ACE-K: acesulfame potassium; CTL, control.

## 4. Discussion

The effects of habitual sugar or LCS consumption during early life periods of development on subsequent energy balance control during adulthood is poorly understood. Here we reveal that early life voluntary consumption of a sucrose solution modeling a concentration and daily caloric levels commonly consumed by humans did not yield any effects on energy intake, body weight, or glucose homeostasis – including any impacts on energy intake and body weight when challenged with a cafeteria-style Western diet (CAF) diet during adulthood. For LCSs, early life habitual ACE-K, but not stevia consumption disrupted energy balance upon adulthood consumption of a CAF diet, with notable sex-dependent effects. Specifically, both male and female ACE-K-exposed rats consumed fewer calories than controls during the adult CAF period, yet the females showed an accompanying significant reduction in body weight whereas the males did not. This sex-dependent discrepancy in ACE-K rats is likely driven by reduced mRNA expression of *BMP8B* and *UCP1* in brown adipose tissue (BAT) in males, whereas no differences were observed in these markers of thermogenic activity in females.

The absence of effects resulting from early life sugar consumption on energy balance parameters was surprising, especially noting the ‘developmental programming hypothesis’, which holds that early life is a critical period for programming energy balance and metabolic health later in life [45]. However, previous research has indicated that adolescent consumption of either *ad libitum* or 5% kcals from sugar of comparable sugar solutions (HFCS or sucrose) in rats did not impart changes in caloric intake or body weight during early adulthood, despite resulting in long-lasting memory impairments and altered microbiome [23,37,46,47]. Present results, together with these previous findings, suggest that early life sugar consumption may have more profound long-term influences on memory function than ingestive and metabolic outcomes.

Previous research has identified sex-specific effects of 4-week adult ACE-K consumption on body weight of mice, such that males had increased body weight compared to controls and females did not differ from controls [48]. These differences were related to changes in gut bacteria community composition and functional genes related to energy metabolism. Our recent work using the early life LCS model revealed no substantial differences in the gut microbiome [49], but this previous study did not include CAF diet consumption during adulthood. Although both our present study and the work by Bian et al. [48] support a dysregulation of energy balance related to ACE-K consumption that is more pronounced in males, there are several important factors to resolve that may be related to the discrepant findings between these two studies: rodent age, duration of exposure to ACE-K (PN 56-84 vs. PN 26-70), dose of ACE-K administered (37.5 mg/kg vs. ~15 mg/kg), route of administration (oral gavage vs. voluntary consumption in supplemental drinking water), and subsequent exposure to a Western/CAF diet. Nevertheless, our current findings provide a clear connection between early life ACE-K exposure within the federal recommended level, Western diet consumption in adulthood, and altered gene expression of markers for thermogenesis, thus revealing an LCS-induced offset in metabolism that is specific to males.

A particularly notable finding from the present work is the effect of early life stevia consumption, with a significant increase in intake of the sugar beverage observed for females during adult CAF diet exposure compared to controls. However, overall caloric intake for stevia-exposed females did not differ from controls, and no differences in any other outcomes were indicated for stevia-exposed males. Previous findings show that high doses of stevia (25, 250, 500, and 1000 mg/kg) result in body weight reductions beginning after 6 weeks of exposure in adult female rats [50], which suggests that a longer period of exposure and higher dose than the ADI of 4 mg/kg may be required in order to observe effects on energy balance for this particular LCS. Similar weight-reducing and notable anti-diabetic properties have been observed with high doses of stevia in diabetic rats [51]. In vitro work has further suggested that stevia may be an endocrine disruptor [52], but this effect has not been documented to date in humans or rodent models, to our knowledge. Regardless, our data indicate that, when restricted within federal recommended daily limits, early life consumption of stevia had no impact on overall energy balance yet increased consumption of a sugary beverage during adulthood in a sex-dependent manner. The mechanisms underlying these results warrant further investigation. We previously revealed that early life LCS (ACE-K) consumption using a similar model as the current study reduced lingual sweet taste receptor expression, and that LCS consumption independent of sweetener (ACE-K, stevia, or saccharin) or sex increased consumption of a sugar solution when offered alongside *ad libitum* healthy chow [49]. These previous findings taken together with the present results collectively indicate that early life LCS consumption appears to have long-lasting effects on voluntary consumption of sugary beverages during adulthood, although it is worth noting that no such outcomes were observed in the present study following ACE-K consumption. Yet, in the present study other food options beyond a sugar solution and healthy chow were presented during the adulthood CAF diet exposure, which could have influenced overall sugar intake.

The distinct effects of early life exposure to ACE-K compared to stevia on adult energy balance parameters are noteworthy, especially considering the varying chemical structures of ACE-K and stevia and their synthetic vs. ‘natural’ origins. ACE-K is rapidly and completely absorbed into the body, yet it is not metabolized prior to being excreted, primarily through urine, within approximately 24 hours after consumption [53–55]. Additionally, it is unknown if, or to what extent, the potassium salt dissociates from the acesulfame anion in ACE-K, and therefore ACE-K consumption could also alter circulating potassium levels, which may affect the normal function of the sodium-potassium pump and ensuing hydrolysis of ATP to generate energy [53,56]. In contrast, stevia (i.e., a mix of steviol glycosides) is more slowly absorbed into the body and is specifically metabolized by colonic bacteria to the common metabolites steviol or steviol glucuronide before being excreted, predominantly in feces [53,57,58]. Although the distinct means of absorption, metabolism, and excretion of ACE-K and stevia are generally comparable in rat models vs. humans, a notable difference is that, in humans, stevia is predominantly excreted in urine instead of feces [53,57]. In light of these differences between ACE-K and stevia, our current findings – that ACE-K affected energy balance and stevia did not – suggest that the rate LCSs are absorbed into the body as well as their metabolism and excretion may be critical distinguishing features in their role on body weight regulation.

Our findings for sex-specific differences in BAT gene expression of markers for thermogenesis during the adult Western diet period for rats after early life ACE-K consumption warrant further investigation. Previous evidence revealed that ACE-K consumption increases insulin and leptin levels [59], which are metabolic hormones with distinct energetic effects in males vs. females [60]. It is possible that slight alterations in insulin and leptin signaling associated with ACE-K consumption could have contributed to the differences in gene expression of markers for thermogenesis observed between males and females in the present study. Although we did not measure insulin and leptin, these may be important systems to examine in future work, especially noting the body of evidence indicating that LCSs can disrupt glucose homeostasis [16,49,61].

Collectively, our findings add further evidence challenging the efficacy of using LCSs for weight management and energy balance control. While early life ACE-K consumption was associated with reduced caloric intake during adulthood, this was accompanied by blunted expression of BAT thermogenic markers that promote energy expenditure, thus resulting in a net zero effect on body weight relative to controls. Importantly, our experiments involved voluntary consumption of LCSs confined within federal recommended daily limits, making the results more readily relevant to humans compared to the bulk of previous rodent LCS research that utilized excessive and involuntary consumption. These results further highlight the importance of sex differences in the impacts of LCS consumption on energy balance, and distinguish early life exposure as a critical period for lasting metabolic effects.

## 5. Conclusions

Our results reveal that habitual consumption of low-calorie sweeteners during early life development influences energy balance outcomes during adulthood, such that ACE-K consumption was linked with blunted expression of genes related to thermogenesis during adulthood in males, and stevia consumption was linked with elevated sugary beverage consumption during adulthood in females.

## Acknowledgment

Fig. 1A, 2A, and 3A graphics were created with BioRender.com.

## Author Contributions

Conceptualization, S.E.K., L.T., A.M.R.H., and L.A.S.; Methodology, S.E.K., L.T., A.M.R.H., and L.A.S.; Formal Analysis, L.T. and A.M.R.H.; Investigation, L.T., A.M.R.H., A.E.K., G.M.S., L.D-S., L.T.L., and M.E.K.; Writing – Original Draft Preparation, A.M.R.H., L.T., and S.E.K.; Writing – Review & Editing, A.M.R.H., L.T., L.A.S., and S.E.K.; Visualization, L.T. and A.M.R.H.; Supervision, S.E.K.; Funding Acquisition, S.E.K.

## Institutional Review Board Statement

This study was conducted according to the guidelines of and approved by the Institutional Animal Care and Use Committee at the University of Southern California.

---

**Supplementary Figure S1:**
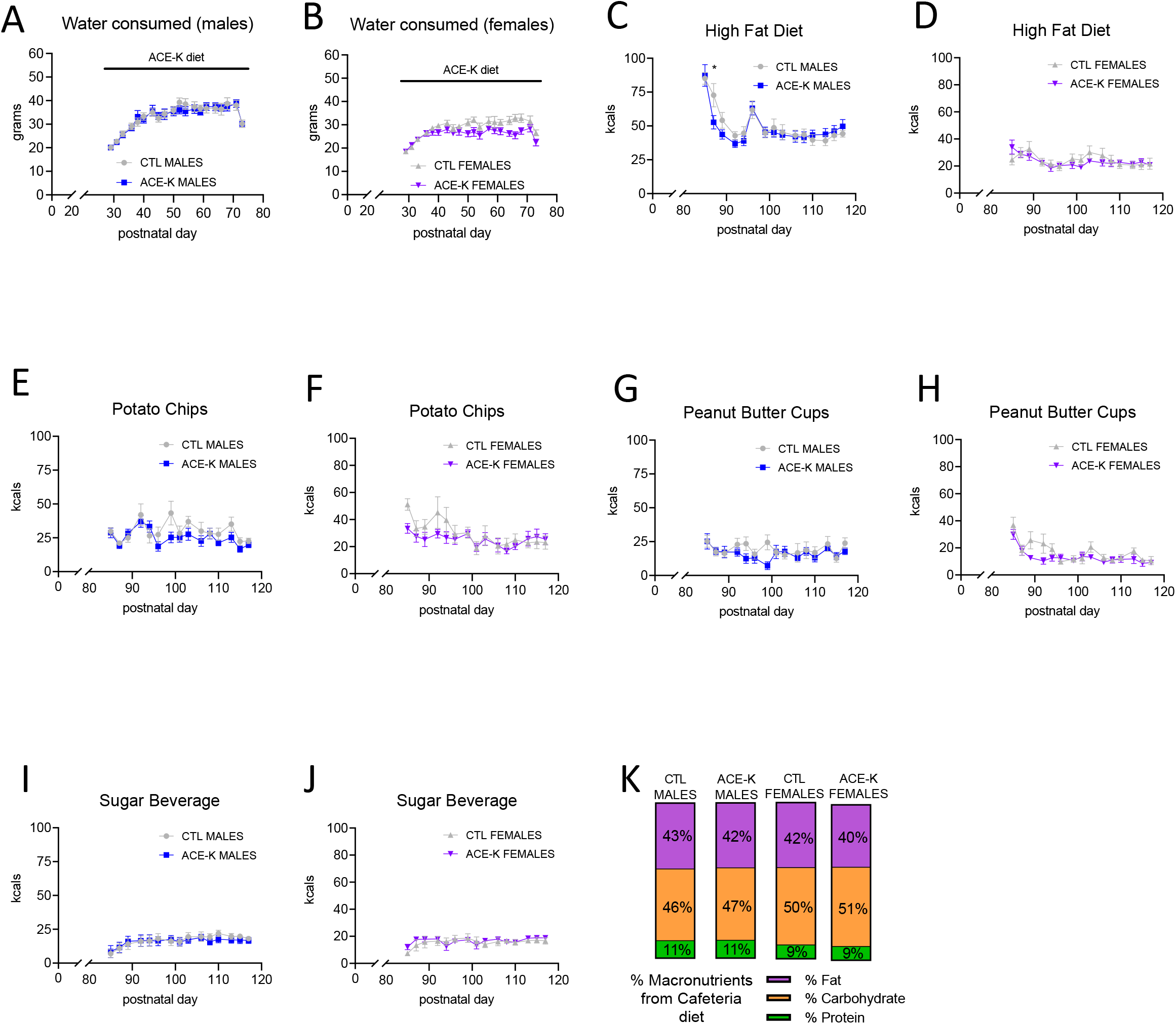
Expanded analyses of water intake and cafeteria diet consumption over time for early life ACE-K-exposed rats. Early life acesulfame potassium (ACE-K) consumption had no effect on water intake in male or female rats (A-B). During the cafeteria (CAF) diet period in adulthood, there was a group *×* time interaction (*p* = 0.02) for total kcals consumed from high fat diet over time in males such that ACE-K males consumed fewer kcals from the high fat diet than control (CTL) males on postnatal day (PN) 84 (C). There were no differences in total kcals consumed over time from any of the other CAF diet components for males (potato chips, peanut butter cups, sugar beverage) and for any of the CAF diet components for females (D-J). Overall percentage of kcals consumed from fat, carbohydrate, and protein during the CAF diet period did not differ between ACE-K and CTL males or females (K). Data are represented as mean ± SEM, \**p* < 0.05; PN: postnatal day; CAF: cafeteria; CTL: control; ACE-K: acesulfame potassium; kcal: kilocalories.

**Supplementary Figure S2:**
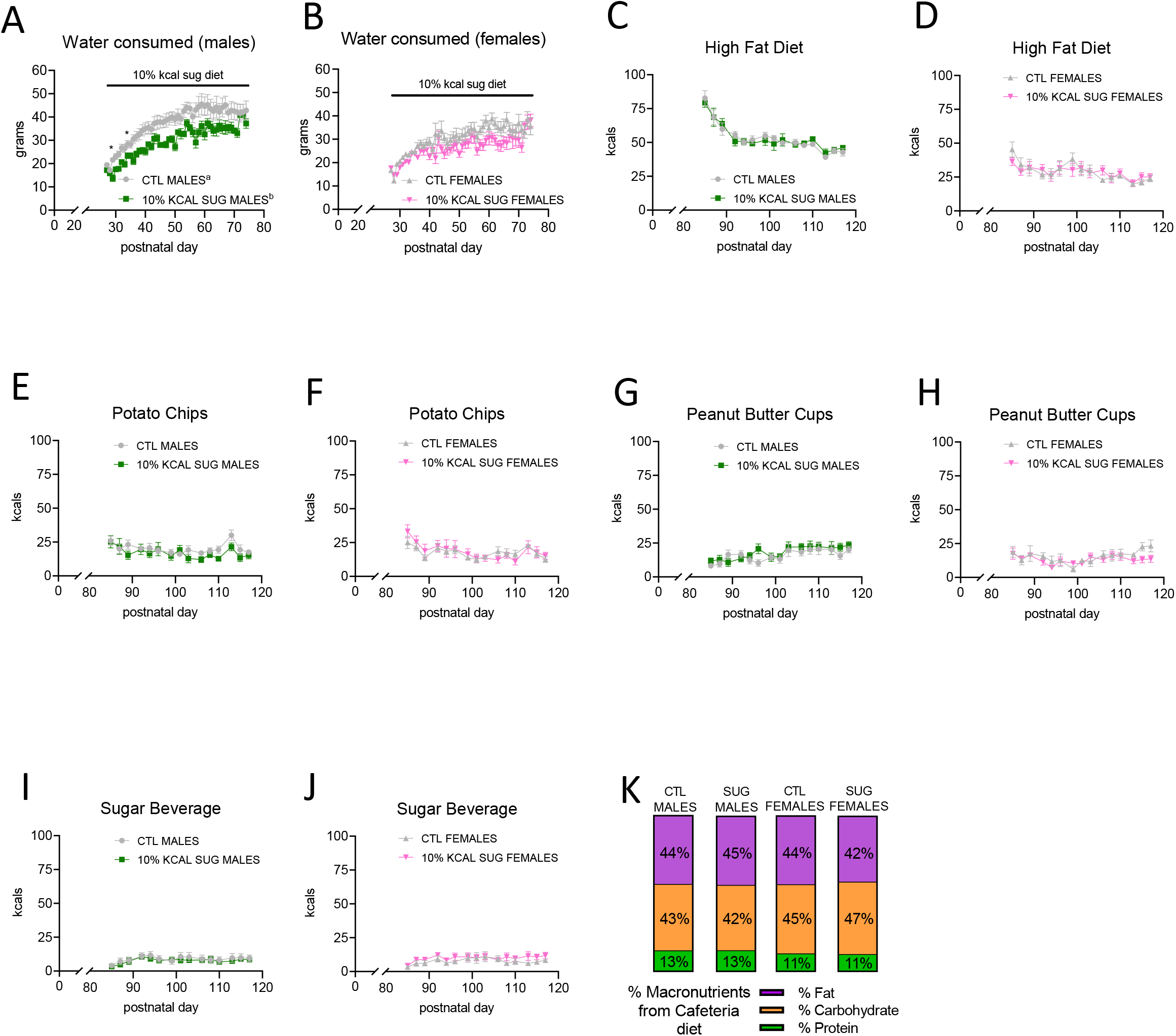
Expanded analyses of water intake and cafeteria diet consumption over time for early life sugar-exposed (10% kcal) rats. Limiting consumption of sugar to 10% of total calories consumed during adolescence (10% KCAL SUG) decreased water intake in male rats (main effect of group [*p* = 0.04] and the group *×* time interaction [*p* = 0.005], with significant post hoc comparisons at PN 29 and 36; A) and female rats (group *×* time interaction [*p* = 0.0004], no significant post hoc comparisons; B). These differences in water intake are likely due to compensation for the volume of sucrose beverage provided per day (10-20 mL). During the cafeteria diet period in adulthood, there were no differences in total kcals consumed over time from any of the cafeteria diet components (high fat diet, potato chips, peanut butter cups, sugar beverage) for males or females (C-J). Overall percentage of kcals consumed from fat, carbohydrate, and protein during the CAF diet period did not differ between 10% KCAL SUG and CTL males or females (K). Data are represented as mean ± SEM, \**p* < 0.05, significant main effect for group indicated by different superscript letters accompanying group names; PN: postnatal day; CAF: cafeteria; CTL: control; SUG: sugar; kcal: kilocalories.

**Supplementary Figure S3:**
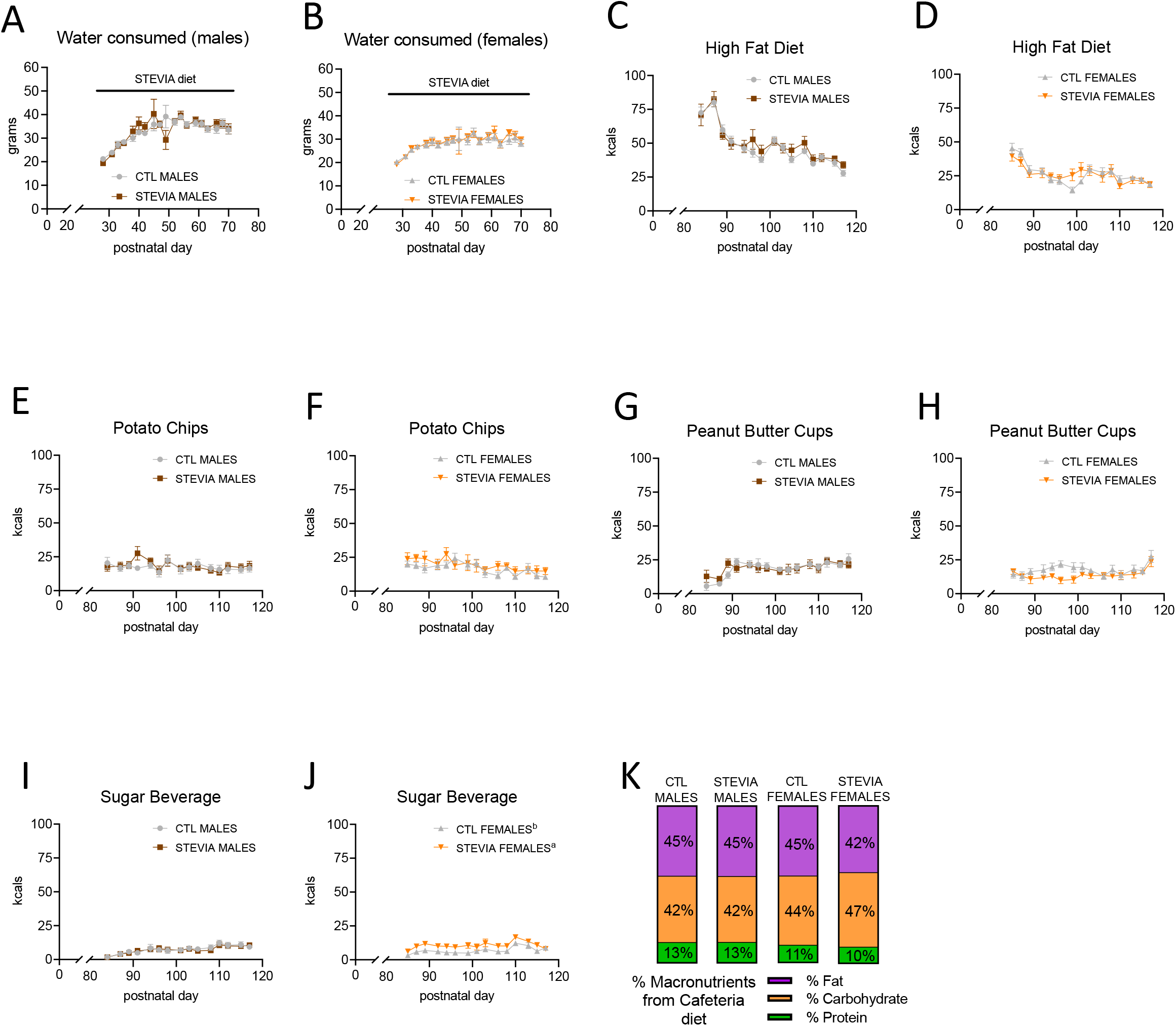
Expanded analyses of water intake and cafeteria diet consumption over time for early life stevia-exposed rats. Early life stevia consumption (STEVIA) had no effect on water intake in male or female rats (A-B). During the cafeteria diet period in adulthood, there were no differences in total kcals consumed over time from the high fat diet, potato chips, or peanut butter cups for males or females (C-H). However, although there was no difference in total kcals consumed over time from the sugar beverage for male STEVIA rats (I), female STEVIA rats consumed more total kcals over time from the sugar beverage than control females (J). Overall percentage of kcals consumed from fat, carbohydrate, and protein during the CAF diet period did not differ between STEVIA and CTL males or females (K). Data are represented as mean ± SEM, \**p* < 0.05, significant main effect for group indicated by different superscript letters accompanying group names; PN: postnatal day; CAF: cafeteria; CTL: control; kcal: kilocalories.

**Supplementary Figure S4:**
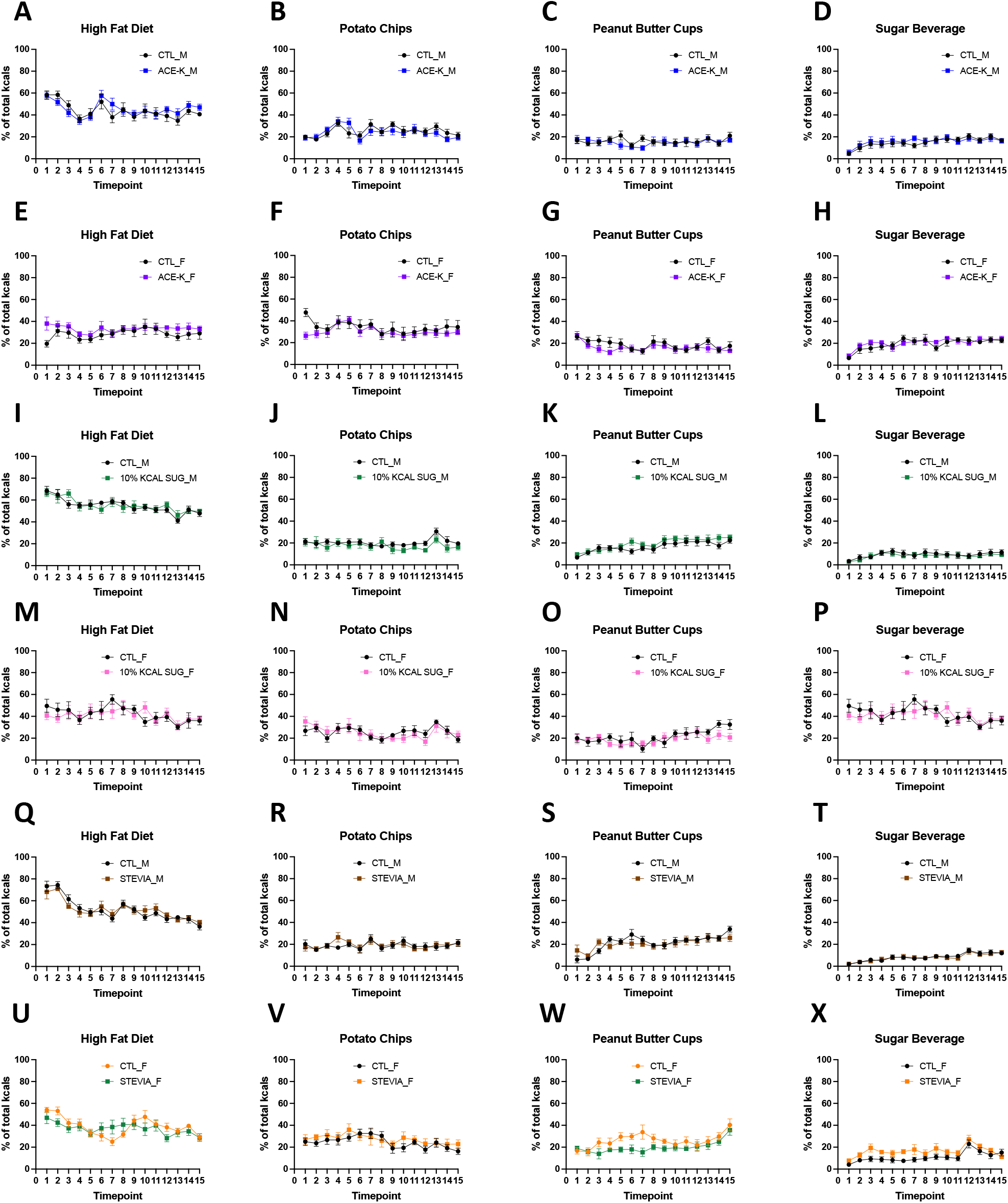
Percentage of total calories consumed from cafeteria diet components during adult exposure. When examined as percentage of total calories consumed from each of the cafeteria diet components (% total kcals from high fat diet, potato chips, peanut butter cups, or sugar beverage), there were no differences for ACE-K males (vs. control males; A-D), ACE-K females (vs. control females; E-H), 10% KCAL SUG males (vs. control males; I-L), 10% KCAL SUG females (vs. control females, M-P), STEVIA males (vs. control males; Q-T), or STEVIA females (vs. control females; U-X). Data are represented as mean ± SEM; PN: postnatal day; CTL: control; kcal: kilocalories.

